# Mannan detecting C-type lectin receptor probes recognise immune epitopes with diverse chemical, spatial and phylogenetic heterogeneity in fungal cell walls

**DOI:** 10.1101/677104

**Authors:** Ingrida Vendele, Janet A. Willment, Lisete M. Silva, Angelina S. Palma, Wengang Chai, Yan Liu, Ten Feizi, Mark H. T. Stappers, Gordon D. Brown, Neil A. R. Gow

## Abstract

During the course of fungal infection, pathogen recognition by the innate immune system is critical to initiate efficient protective immune responses. The primary event that triggers immune responses is the binding of Pattern Recognition Receptors (PRRs), which are expressed at the surface of host immune cells, to Pathogen-Associated Molecular Patterns (PAMPs) located predominantly in the fungal cell wall. Most fungi have mannosylated PAMPs in their cell walls and these are recognized by a range of C-type lectin receptors (CTLs). However, the precise spatial distribution of the ligands that induce immune responses within the cell walls of fungi are not well defined. We used recombinant IgG Fc-CTLs fusions of three murine mannan detecting CTLs, including dectin-2, the mannose receptor (MR) carbohydrate recognition domains (CRDs) 4-7 (CRD4-7), and human DC-SIGN (hDC-SIGN) and the β-1,3 glucan-binding lectin dectin-1 to map PRR ligands in the fungal cell wall. We show that epitopes of mannan-specific CTL receptors can be clustered or diffuse, superficial or buried in the inner cell wall. We demonstrate that PRR ligands do not correlate well with phylogenetic relationships between fungi, and that Fc-lectin binding discriminated between mannosides expressed on different cell morphologies of the same fungus. We also demonstrate CTL epitope differentiation during different phases of the growth cycle of *Candida albicans* and that MR and DC-SIGN labelled outer chain *N*-mannans whilst dectin-2 labelled core *N*-mannans displayed deeper in the cell wall. These immune receptor maps of fungal walls therefore reveal remarkable spatial, temporal and chemical diversity, indicating that the triggering of immune recognition events originates from multiple physical origins at the fungal cell surface.

**Author Summary:** Invasive fungal infections remain an important health problem in immunocompromised patients. Immune recognition of fungal pathogens involves binding of specific cell wall components by pathogen recognition receptors (PRRs) and subsequent activation of immune defences. Some cell wall components are conserved among fungal species while other components are species-specific and phenotypically diverse. The fungal cell wall is dynamic and capable of changing its composition and organization when adapting to different growth niches and environmental stresses. Differences in the composition of the cell wall lead to differential immune recognition by the host. Understanding how changes in the cell wall composition affect recognition by PRRs is likely to be of major diagnostic and clinical relevance. Here we address this fundamental question using four soluble immune receptor-probes which recognize mannans and β-glucan in the cell wall. We use this novel methodology to demonstrate that mannan epitopes are differentially distributed in the inner and outer layers of fungal cell wall in a clustered or diffuse manner. Immune reactivity of fungal cell surfaces did not correlate with relatedness of different fungal species, and mannan-detecting receptor-probes discriminated between cell surface mannans generated by the same fungus growing under different conditions. These studies demonstrate that mannan-epitopes on fungal cell surfaces are differentially distributed within and between the cell walls of fungal pathogens.

## Introduction

Fungi are associated with a wide spectrum of diseases ranging from superficial skin and mucosal surface infections in immunocompetent people, to life-threatening systemic infections in immunocompromised patients [1, 2]. The global burden of fungal infections has increased due to infection related or medically imposed immunosuppression, the use of broad-spectrum antibiotics that suppress bacterial competitors, and the use of prosthetic devices and intravenous catheters in medical treatments [3, 4]. Patients that are pre-disposed to fungal diseases include those with neutropenia, those undergoing stem cell or organ transplant surgery or recovering from surgical trauma as well as HIV infected individuals and those with certain rare predisposing mutations in immune recognition pathways [3–6].

Innate immunity is the primary defence mechanism against fungal infections and involves host Pattern Recognition Receptors (PRRs) that recognise specific Pathogen-Associated Molecular Patterns (PAMPs), which are mostly located within the cell wall [7–9]. These receptor-ligand interactions are the primary origin of all immune responses and they promote expression and secretion of various chemokines and cytokines that results in recruitment of neutrophils, macrophages and other immune cell types to the site of infection, which ultimately leads to containment and clearance of the pathogen and the activation of protective longer term adaptive immunity [9–12].

The repertoire of ligands in fungal cell walls that engage with cognate PRRs has been reviewed extensively [13–15] but we lack information about where precisely these ligands are located in the fungal cell wall. Most fungi have a two layered cell wall with an inner layer comprised of a conserved glucan-chitin scaffold to which a diversity of outer cell wall components are attached that varies significantly between different fungal species [14–16]. In *Candida* species the outer wall is dominated by a fibrillar layer of highly glycosylated cell wall proteins that are extensively decorated with *N*- and *O*-linked mannans and phosphomannans [17, 18]. The chemical fingerprint of the fungal cell wall mannans are used to identify medically relevant species in diagnostic tests [19–21]. However, within a species the composition of the cell wall is highly variable and changes according to morphology, growth stage, nutrient availability, the presence of antibiotics and other environmental stressors [15, 22–25]. This chemical and architectural plasticity represents a moving target for the immune system and leads to differential immune activation at different stages of an infection [25]. Understanding the relationship between fungal cell wall composition and immune recognition is therefore critically important in fungal pathogenesis and immunity and in the context of fungal diagnostics, vaccines and immunotherapies [26–28].

C-type Lectin (CTL) receptors orchestrate antifungal immunity through recognition of fungal-specific ligands that are mainly located in the cell wall [29–33]. Multiple CTLs participate in pathogen recognition of fungal cell wall components including β-glucan, chitin, mannans and melanin [34–40]. Dectin-1 recognises β-1,3-glucan that is a conserved element of the inner cell wall of all known fungal pathogens [41, 42]. Mannans are more complex, comprising linear and branched polymers of mannose sugars linked via α-1,2, α-1,3, α-1,4, α-1,6, and β-1,2 glycosidic bonds that may be further modified by phosphodiester side chains [18, 43, 44]. These glycosides decorate the cell wall proteins of the outer cell walls and the integral proteins in the cell membrane and may account for more than 80% of the mass of the glycoprotein [18, 43, 44]. A wide range of mannan-recognising immune receptors are present in myeloid and epithelial cells including the mannose receptor (MR), dectin-2, dectin-3, mincle, DC-SIGN, galectin-3, FcγR, CD14, CD23, TLR2, TLR4 and TLR6 [13, 34, 37, 45–54]. The number and diversity of mannan-recognising PRRs underlines their importance in primary immune recognition events. Although these recognition events are the primary trigger of the immune response, the precise chemical nature and location of the ligands that are recognised by these immune receptors has not been investigated in detail.

We utilized murine CTL receptor carbohydrate recognition domains and human IgG Fc fusion proteins (Fc-lectins) including dectin-2, MR CRD4-7, dectin-1 and acquired commercial human DC-SIGN-Fc to explore the distribution of mannan- and β-glucan-detecting CTLs [36, 37, 55]. We used the murine MR cysteine rich (CR) domain fused to human IgG Fc to control for Fc mediated binding events [56, 57]. The CR lacks the carbohydrate recognition domains and binds to untreated sulphated carbohydrates. These Fc-lectin probes were used to examine the distribution of immune epitopes in fungal cell surfaces and to examine the nature of the ligand engaging with specific mannan-detecting CTLs. We reveal remarkable diversity in the location and distribution of the cognate immune ligands and demonstrate that these ligands bind different mannans in different parts of the cell wall in order to induce immune responses.

## Results

### Distribution of C-type lectin receptor ligands on fungal cell surfaces

We first confirmed the ability of a set of Fc-lectins to bind their cognate target antigens by ELISA using whole yeast *C. albicans* cells, purified S. *cerevisiae* mannan and *Candida* yeast β-glucan (S Fig. 1 A) and verified the molecular weight of the recombinant Fc-lectins (S Fig. 1 B). The Fc region did not influence binding since the control protein CR-Fc lacking the MR carbohydrate binding domains did not recognise any of the immobilised targets (S Fig. 1 A). Binding of the various Fc-lectins against reported mannan or β-1,3 glucan targets and Fc-probe integrity was therefore confirmed.

We then examined variability in expression and distribution of epitopes for CTL receptors in a range of fungal strains and species (Fig. 1). CTL Fc-lectins bound different yeast cells with differing intensities and patterns (Fig. 1 A-C). MR probe (CRD4-7-Fc) and dectin-2-Fc labelled *C. glabrata* and *C. krusei* more strongly than *C. parapsilosis* (Fig. 1 A, B), however there was no clear correlation between the profiles of Fc-lectin binding and the phylogenetic relatedness of the species. For example, *C. albicans* and *C. dubliniensis* are relatively close genetic relatives yet displayed very different CTL binding profiles (Fig. 1 A, B). *C. dubliniensis* demonstrated higher binding by dectin-2-Fc and CRD4-7-Fc compared to *C. albicans* (Fig. 1 A, B), and mannose receptor probe (CRD4-7-Fc) bound *C. dubliniensis* more diffusely whilst dectin-2-Fc binding was not visible by fluorescence microscopy on *C. albicans* yeast cell walls but labelled *C. dubliniensis* yeast cells in a punctate pattern (Fig. 1 C).

**Figure 1.**
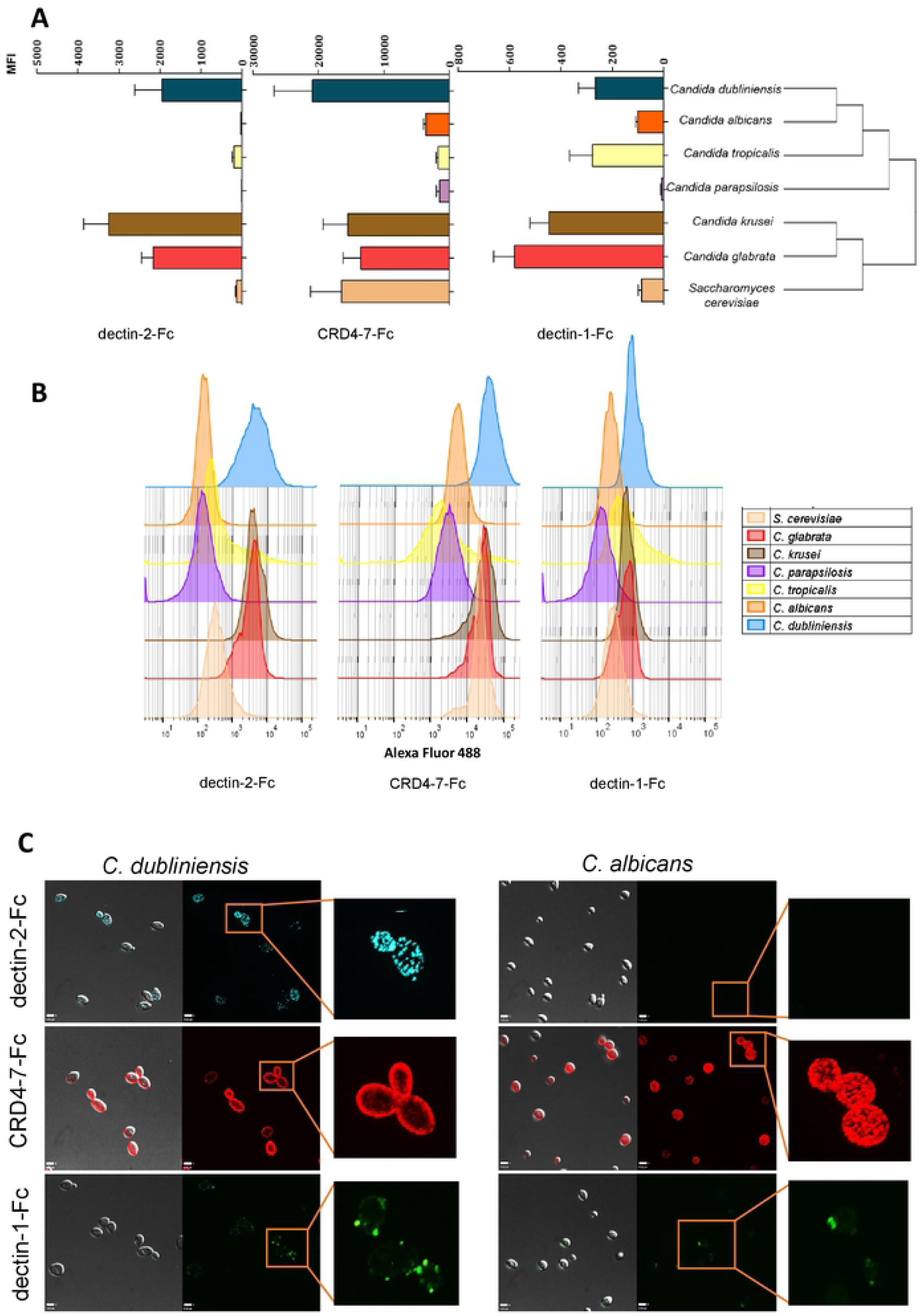
Binding of CTL receptors does not correlate with fungal phylogenetic relationships. (A) Fc-lectin binding to different *Candida* species and *S. cerevisiae* displayed according to phylogenetic relationships in agreement with the published dendrogram [94] and represented as Median Fluorescence Intensity (MFI) of probe binding. (B) Representative images of FACS histograms. (C) Comparison of Fc-lectin binding pattern to evolutionary closely related *C. albicans* and *C. dubliniensis* yeast cells. For immunofluorescence staining and flow cytometry experiments, 2.5 × 10^6^ cells were used in each analysis. Data were obtained using a BD Fortessa flow cytometer and median fluorescence values were used for quantification of probe binding. 3D visualisation on an UltraView® VoX spinning disk confocal microscope was also used to observe binding patterns of Fc-lectins on the cell surfaces. Represented data are means ± SEM, n=3. Scale bars represent 4 µm.

Three virulent (SC5314, Ysu751, J990102) and three attenuated (IHEM3742, AM2003/0069, HUN92) isolates of *C. albicans* were examined, as determined in both mouse and insect systemic models of infection [58, 59]. There was no clear relationship between binding of dectin-2-Fc, CRD4-7-Fc and dectin-1-Fc probes and virulence except that attenuated yeasts exhibited a tendency to higher binding of dectin-2-Fc (Fig. 2 A). However, strain CAI4-CIp10 [60], which is the genetic background for the generation of multiple *C. albicans* null mutants and its progenitor clinical isolate SC5314 exhibited identical Fc-lectin binding patterns (Fig. 2 A). In *C. albicans* hyphae, similar Fc-lectin binding patterns were observed for all strains, however, the attenuated HUN92 isolate was a high dectin-2-Fc binder with most labelling occurring at the hyphal tips (Fig. 2 B). Therefore, the Fc-lectins demonstrated differing binding profiles to different *Candida* species and *C. albicans* strains with no clear association between Fc-lectin binding, phylogenetic relatedness and relative virulence.

**Figure 2.**
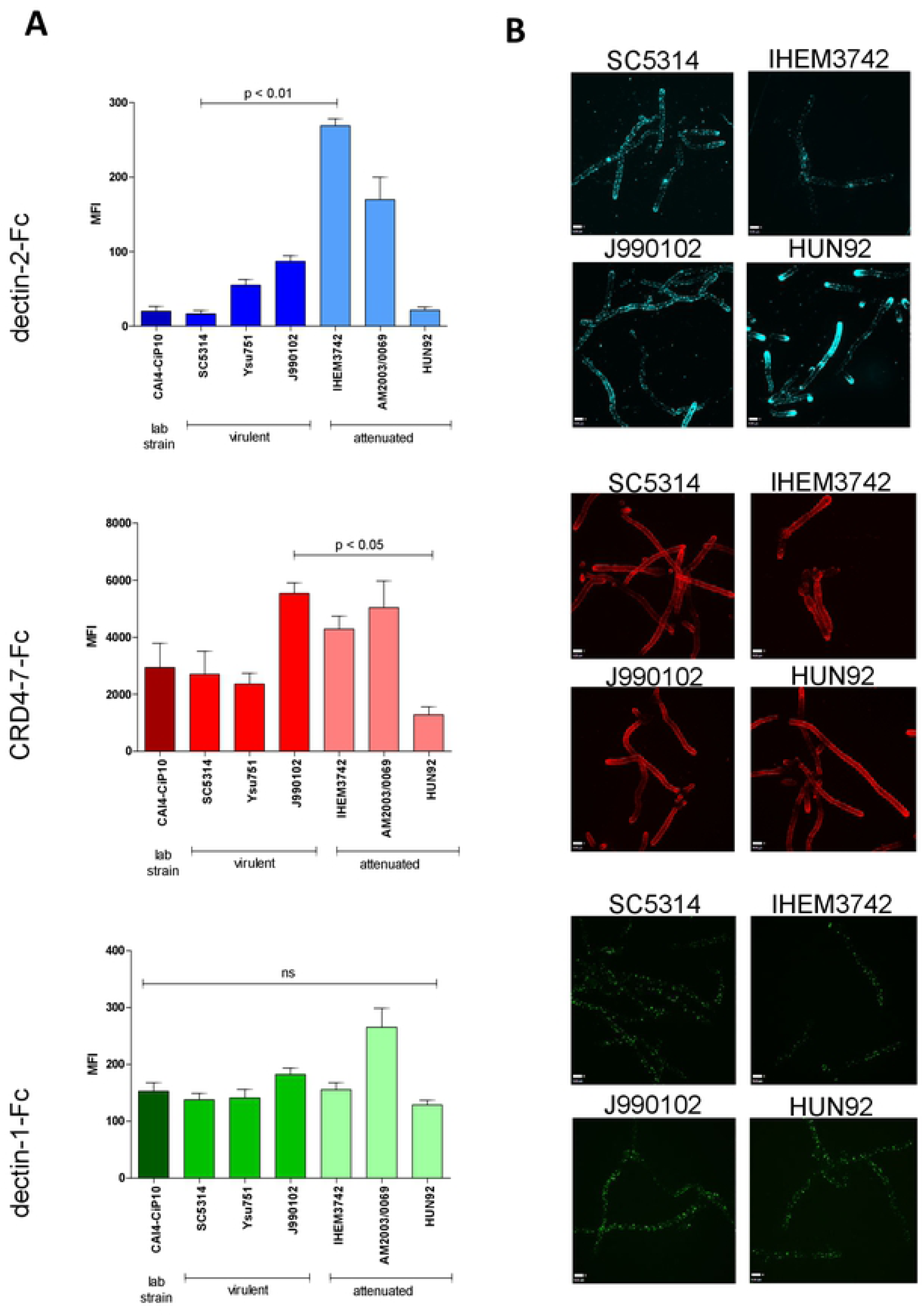
Binding of Fc-lectin probes to different *C. albicans* isolates. (A) Binding of Fc-lectin probes to the laboratory control strain (CAI4-CIp10), and to virulent (SC5314, Ysu751, J990102) and attenuated (IHEM3742, AM2003/0609, HUN92) clinical isolates. Binding to yeast cells is represented as Median Fluorescent Intensities (MFI). (B) Indirect immunofluorescence staining of hyphae of virulent *C. albicans* (SC5314, J990102) and attenuated (IHEM3742, HUN92) isolates with Fc-lectin probes. For immunofluorescence staining and flow cytometry experiments, 2.5 × 10^6^ cells were used per analysis. Data were obtained using a BD Fortessa flow cytometer and 3D visualisation on an UltraView® VoX spinning disk confocal microscope was also used to observe Fc-lectin binding patterns. Median fluorescence values were used for quantification and results are represented as means ± SEM, n=3. Statistical analyses were performed using Kruskal-Wallis with Dunn’s post-hoc test. Scale bars represent 4 µm.

### Differential expression of ligands for C-type lectin receptors during growth and morphogenesis

We next tested the stability of the epitopes recognised by CTL-Fc-lectins during the growth and morphogenesis of *C. albicans*. Samples in the lag phase, early, mid and late exponential as well as stationary phases were sampled during batch growth (Fig. 3 A). During the period of exponential growth of yeast cells, dectin-2-Fc ligand exposure was somewhat reduced (Fig. 3 B) whilst β-glucan for dectin-1-Fc was increased (Fig. 3 B). In contrast, epitopes for mannose receptor assessed by CRD4-7-Fc binding appeared to be exposed throughout all growth phases of batch growth (Fig. 3 B). Therefore PAMP binding was affected by growth phase and potentially growth rate of the target pathogen.

**Figure 3.**
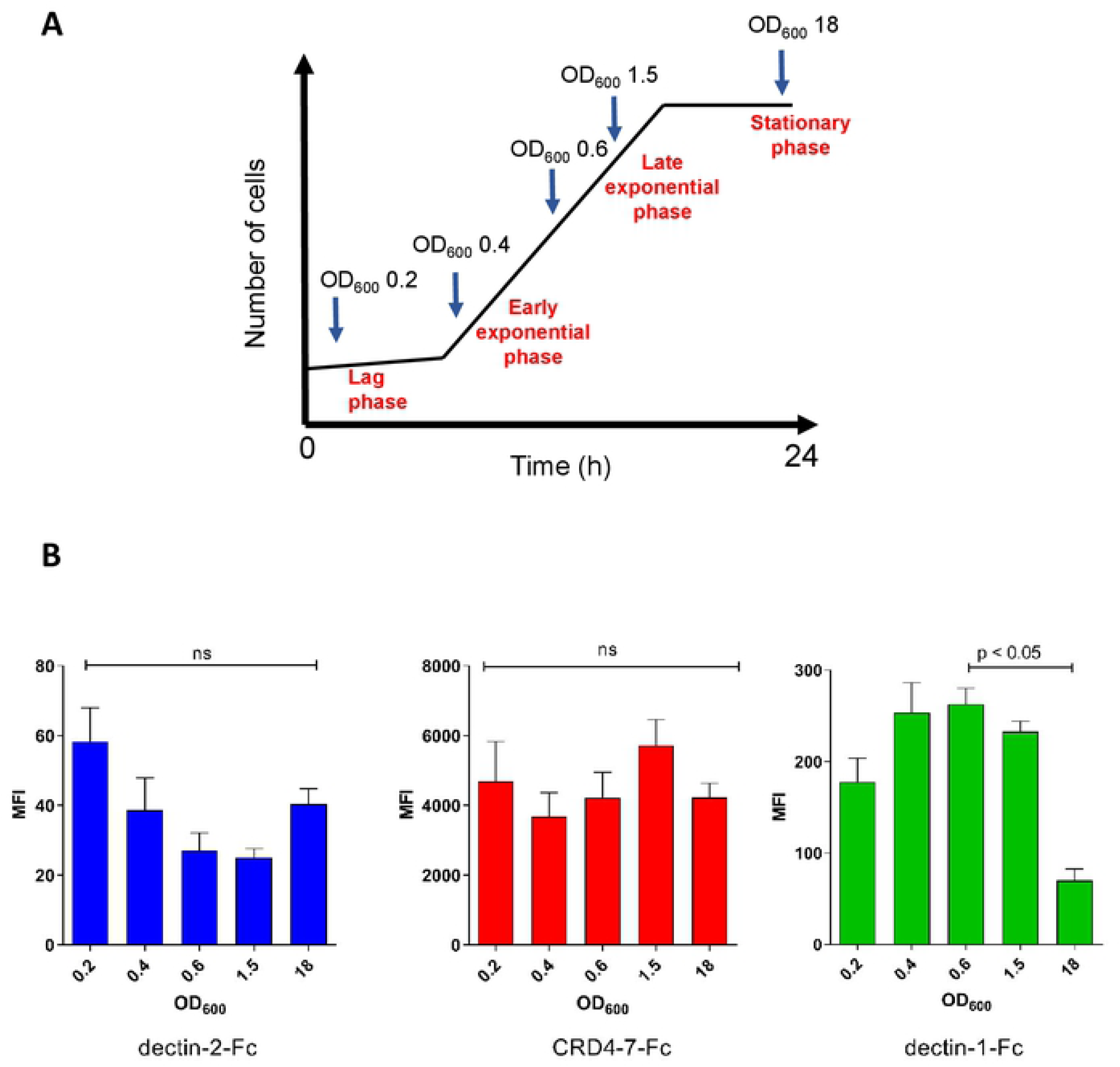
Expression of ligands for CTL receptors during the batch growth of ***C. albicans*.** (A) The OD_600_ values of samples taken for analysis during different growth stages are indicated. (B) Median Fluorescent Intensity (MFI) values from indirect immunofluorescence staining of *C. albicans* (SC5314) yeast cells by dectin-2-Fc, CRD4-7-Fc and dectin-1-Fc. For immunofluorescence staining and flow cytometry experiments, 2.5 × 10^6^ cells were used per analysis. Data were obtained using a BD Fortessa flow cytometer and median fluorescence values were used for quantification. Results are represented as means ± SEM, n=3. Statistical analysis were performed using Kruskal-Wallis with Dunn’s post-hoc tests.

*C. albicans* filamentation was induced and hyphal cells were fixed at different time points to test Fc-lectin binding (Fig. 4). Mannan-recognising dectin-2-Fc and CRD4-7-Fc demonstrated strong binding to early germ tubes grown in serum-containing medium (Fig. 4 A, B). However, binding of both mannan-detecting lectin probes gradually decreased over prolonged periods of hyphal growth (Fig. 4 A, B). In particular, decreasing binding of CRD4-7-Fc to the mother yeast cell was observed and was virtually absent on germ tubes that were older than 2 h (Fig. 4 B). In contrast, although germ tubes lacked bud scars, which have exposed β-1,3 glucan, dectin-1-Fc demonstrated the opposite pattern with binding gradually increasing to the lateral cell walls of maturing filamentous cells (Fig. 4 C). These results reinforce previous observations that nascent mannan epitopes are gradually modified as the yeast cells and hyphae progress through different growth stages [61].

**Figure 4.**
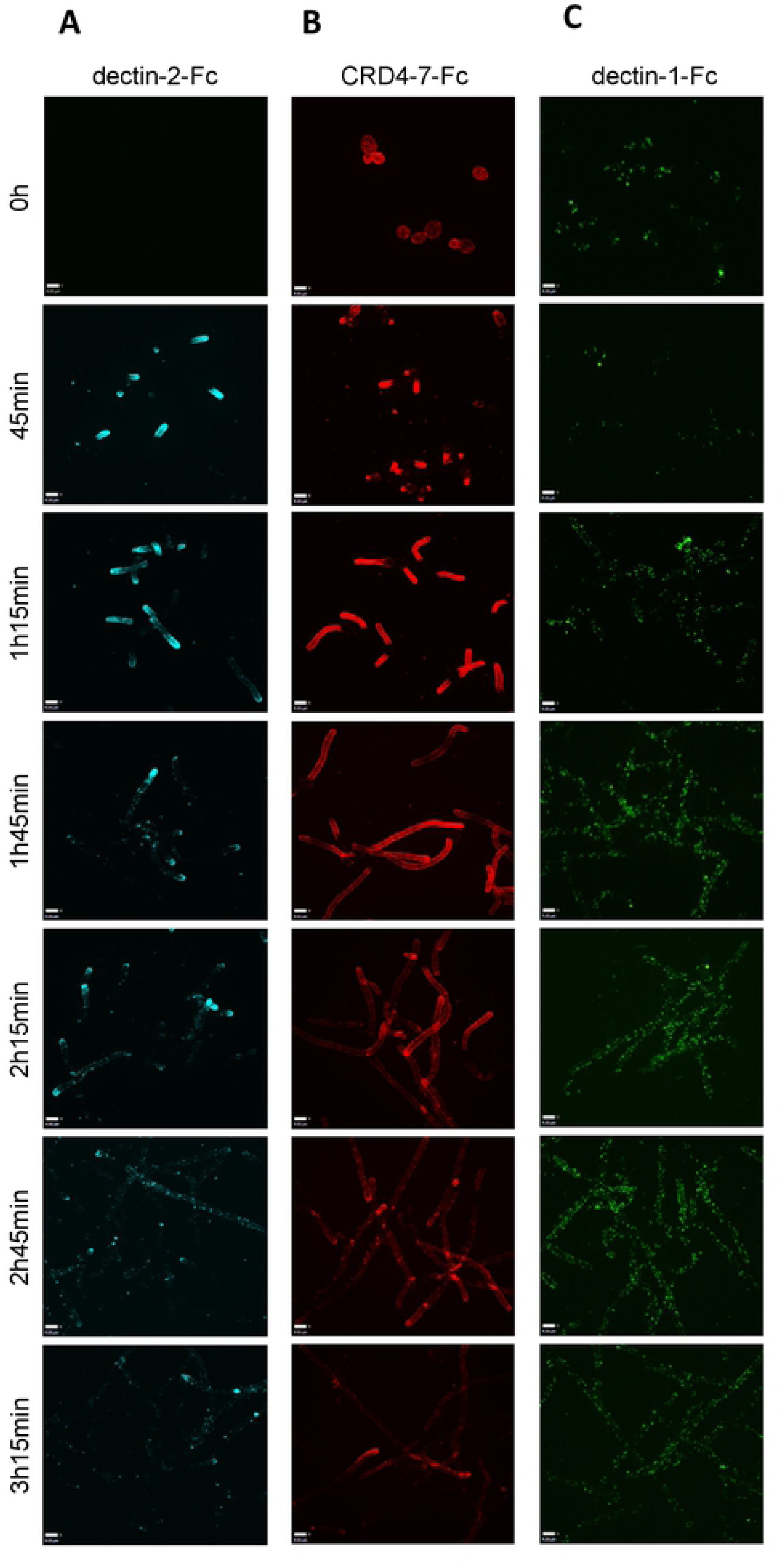
Indirect immunofluorescence of Fc-lectin binding to target ligands in the. *C. albicans* cell wall during different stages of hyphal growth. Representative images of immunofluorescence staining of Fc-lectin binding to *C. albicans* (SC5314) hyphae over prolonged periods of growth by dectin-2-Fc (A), CRD4-7-Fc (B) and dectin-1-Fc (C). Data were obtained using UltraView® VoX confocal spinning disk microscope and scale bars represent 4 µm.

The binding of mannan-detecting C-type lectins to yeast, pseudohypha, hypha and the recently described goliath cells [62] of C*. albicans* was examined (Fig. 5). Dectin-2-Fc demonstrated low binding affinity to *C. albicans* fixed yeast cells which could be detected by flow cytometry (Fig. 1–3) but not by microscopy (Fig. 5 A). Fixation was used to capture and immobilise cells at specific morphogenetic stages, but control experiments showed that paraformaldehyde fixation did not influence Fc-lectin binding patterns (data not shown). Dectin-2-Fc exhibited punctate binding pattern on hyphae with strong staining observed at the hyphal tip (Fig. 5 A). In contrast, CRD4-7-Fc demonstrated high intensity punctate binding to both yeast cells and hyphae (Fig. 5 B). As predicted, dectin-1-Fc recognised yeast cells mainly at the bud scars while some punctate binding was also detected along hyphae (Fig. 5 C) [63]. All Fc-lectins recognised pseudohypha cells with intermediate binding strengths compared to that for yeast and hyphae (Fig. 5 A-C). Recently, goliath cells have been observed as a form of cellular gigantism in *Candida* species [59]. Fc-lectin binding to *C. albicans* goliath cells revealed punctate dectin-2-Fc binding while CRD4-7-Fc bound more uniformly around the cell surface (Fig. 5). As before, dectin-1-Fc bound mainly to the bud scars of goliath cells (Fig. 5). Negative control protein CR-Fc did not show any binding to any of the *C. albicans* morphologies (Fig. 5 D), and binding to yeast cells was not detected by flow-cytometric analyses. These data demonstrated differences in the specificities of dectin-2-Fc and CRD4-7-Fc towards the fungal cell surface components, and were in accord with knowledge of the glycan-binding specificities of dectin-1 [64] and of CR-Fc [57].

**Figure 5.**
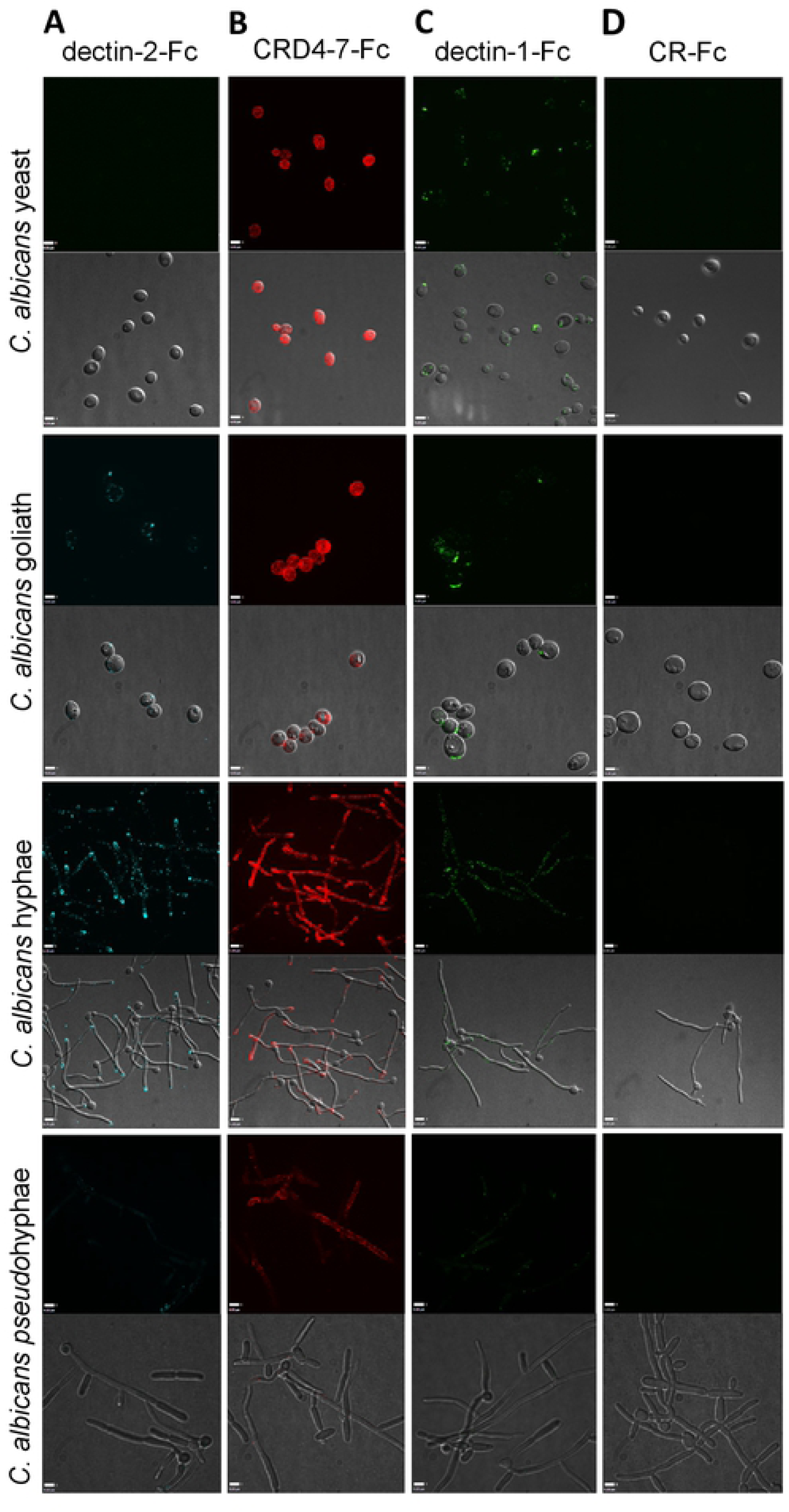
Indirect immunofluorescence of Fc-lectin binding to different *C. albicans* morphological forms. Representative images of immunofluorescence staining of Fc-lectin binding to *C. albicans* yeast, hyphae, pseudohyphae (SC5314) and goliath (BWP17 + Clp30) cells by dectin-2-Fc (A), CRD4-7-Fc (B), dectin-1-Fc (C) and CR-Fc (D). Data were obtained using UltraView® VoX confocal spinning disk microscope and scale bars represent 4 µm for yeast and pseudohyphae images and 6 µm for hyphae.

### Spatial distribution of mannan epitopes in the inner cell wall

To elucidate precise localisation of ligands for mannan-recognising Fc-lectins within the cell wall, immunogold labelling of dectin-2-Fc, CRD4-7-Fc and CR-Fc-stained embedded sections of cells was analysed by TEM (Fig. 6). We observed clustered dectin-2-Fc binding to both yeast and hyphae inner cell walls of *C. albicans* with little labelling of the outer mannoprotein-rich fibrils (Fig. 6 A). CRD4-7-Fc recognised ligands within the plasma membrane as well as outer glycoprotein fibrils (Fig. 6 B). CR-Fc gave no staining (Fig. 6 C). The differential specificities of dectin-2-Fc, CRD4-7-Fc and CR-Fc recognition were again demonstrated.

**Figure 6.**
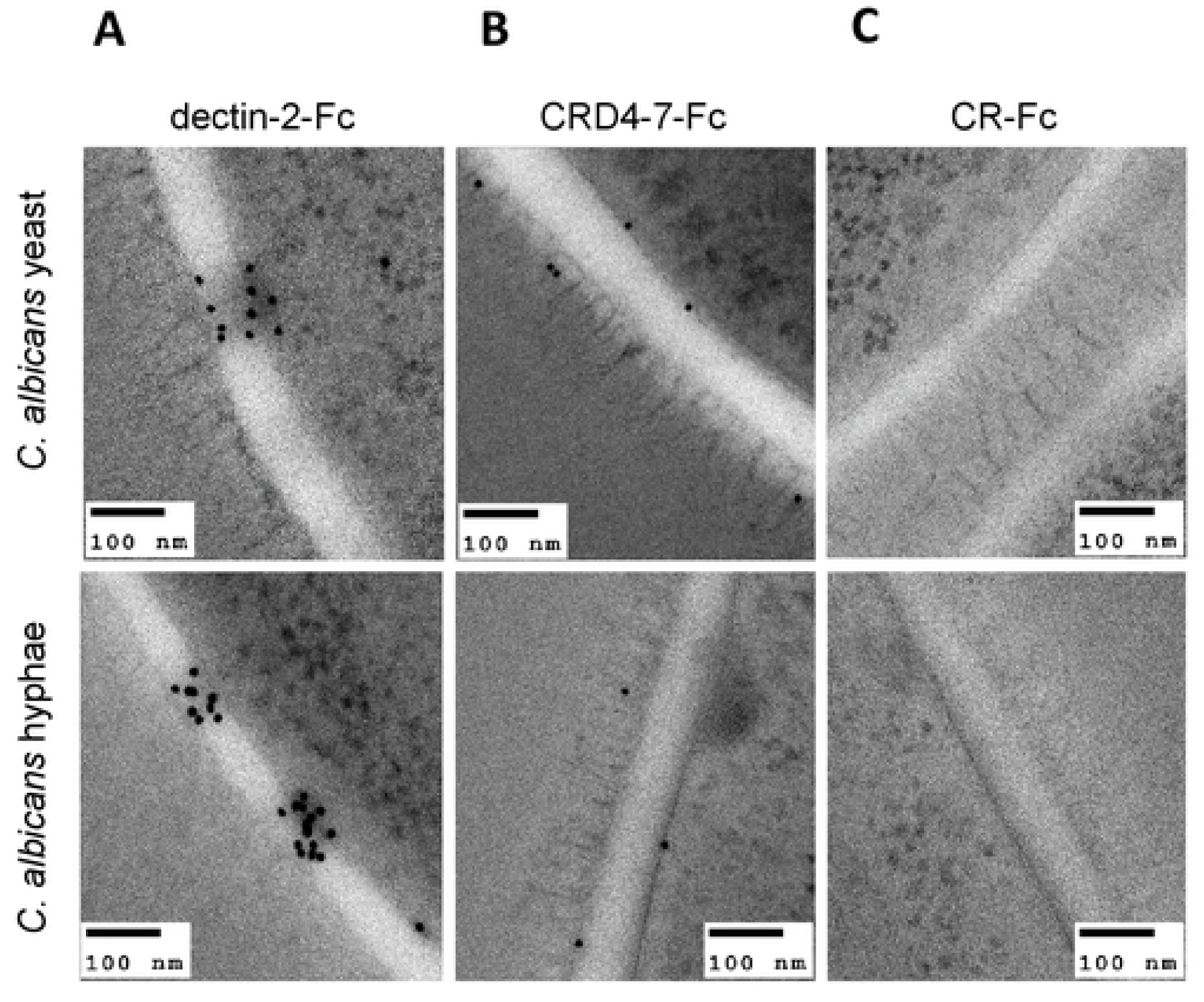
Immunogold localisation of Fc-lectin probes on *C. albicans* yeast and hypha cell walls. Representative TEM images of at least one experiment where Protein A gold conjugate was used to detect dectin-2-Fc (A), CRD4-7-Fc (B) and CR-Fc (C) binding on *C. albicans* (SC5314) cell walls. TEM images were taken using a JEM-1400 Plus using an AMT UltraVUE camera. Scale bars represent 100 nm.

Flow cytometry and microscopy were used to compare binding strengths of Fc-lectins to *C. albicans* yeast cell wall after mild heat treatment (at 65°C) which mechanically perturbs the normal cell wall architecture resulting in the permeabilising of the wall to otherwise impermeable high molecular weight components (Fig. 7). This mild heat treatment is often used to heat-kill (HK) cells to prevent cellular morphogenesis during immunological examinations. Binding affinities of dectin-2-Fc, CRD4-7-Fc and dectin-1-Fc to formaldehyde-fixed or HK *C. albicans* yeast cells were compared (Fig. 7 A-C). Dectin-2-Fc binding increased significantly following HK treatment (Fig. 7 A) while CRD4-7-Fc binding was reduced to a minor extent (Fig. 7 B), suggesting that the CRD4-7-Fc MR ligand was superficial whilst the dectin-2 ligand was buried deeper in the cell wall and was initially inaccessible to CTLs. HK also increased binding of dectin-1-Fc, due to β-glucan exposure (Fig. 7 C). The specificity of Fc-lectin binding was further corroborated by blocking the binding of the Fc-lectins with purified *C. albicans* cell wall mannan (dectin-2-Fc and CRD4-7-Fc) or yeast cell wall β-glucan (dectin-1-Fc) (Fig. 7 A-C). In all cases external addition of an excess of the target polysaccharide completely blocked the binding of Fc-lectins to the cell wall. CR-Fc included as a negative control again showed that binding to fixed or HK cells was not mediated by the Fc region of the CTL-probes (Fig. 7 A-C). Collectively these analyses revealed that some CTL epitopes were clustered in the cell wall whilst others were uniformly distributed and some were exposed and some cloaked within the inner cell wall.

**Figure 7.**
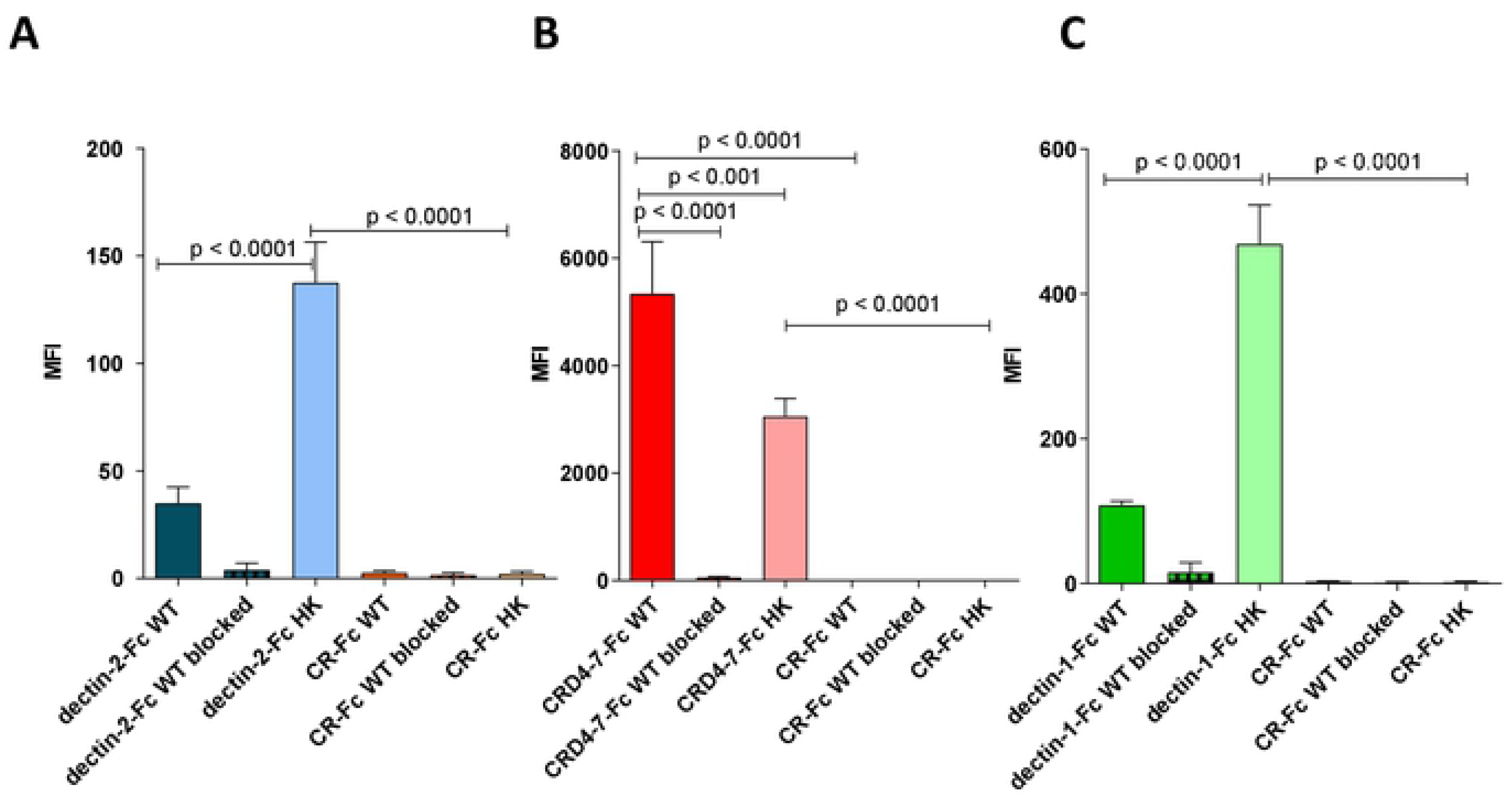
Binding of Fc-lectin probes to *C. albicans* fixed and heat-killed yeast cells. Quantification of Fc-lectin probe binding to WT (live) and HK (heat-killed) cells for dectin-2-Fc (blue) (A), CRD4-7-Fc (red) (B), dectin-1-Fc (green) (C) and CR-Fc (orange) (A-C) binding to WT *C. albicans* yeast cells (dark bars), HK *C. albicans* yeast cells (light bars). Blocking experiments are also shown in which Fc-lectins were preincubated with purified *C. albicans* cell wall mannan (25 µg/ml) or β-glucan (for dectin-1-Fc, 100 µg/ml) and subsequent binding to WT 4% paraformaldehyde fixed *C. albicans* yeast cells (crossed bars). For flow cytometry experiments, 2.5 × 10^6^ cells were used per analysis. Data were obtained using a BD Fortessa flow cytometer and median fluorescence values were used for quantification of probe binding. Data are means ± SEM, n = 3. Statistical analyses with One Way Anova with Tukey’s post-hoc test.

### hDC-SIGN epitopes in the plasma membrane and cell wall

Human DC-SIGN-Fc recombinant protein (Life Technologies) was used to further investigate mannan epitope variability on the fungal surface. hDC-SIGN-Fc consistently demonstrated high affinity binding of *C. albicans* fixed yeast cells (Fig. 8 A) whereas with HK yeast cells there was a significant decrease in hDC-SIGN-Fc binding (Fig. 8 A). Binding was completely blocked in the presence of purified and soluble *C. albicans* mannan (Fig. 8 A, B) and decreased by 75% when using *S. cerevisiae* mannan (Fig. 8 B). Microscopy revealed a high intensity punctate binding pattern on yeast, hyphae and goliath cells and slightly less to pseudohyphal *C. albicans* cells (Fig. 8 C). Using immunogold-labelling hDC-SIGN-Fc epitopes were observed in the plasma membrane, inner cell wall and outer fibrils for yeast and hyphae of *C. albicans* (Fig. 8 D) that was similar to that observed for CRD4-7-Fc labelling. The binding to hDC-SIGN-Fc epitopes progressively decreased in maturing, more elongated *C. albicans* hyphae (Fig. 8 E). Reminiscent of CRD4-7-Fc binding, hDC-SIGN-Fc demonstrated high intensity binding to *C. albicans* cells in different morphologies with binding sites distributed along the plane of the plasma membrane and in the outer cell wall glycosylated fibril layer.

**Figure 8.**
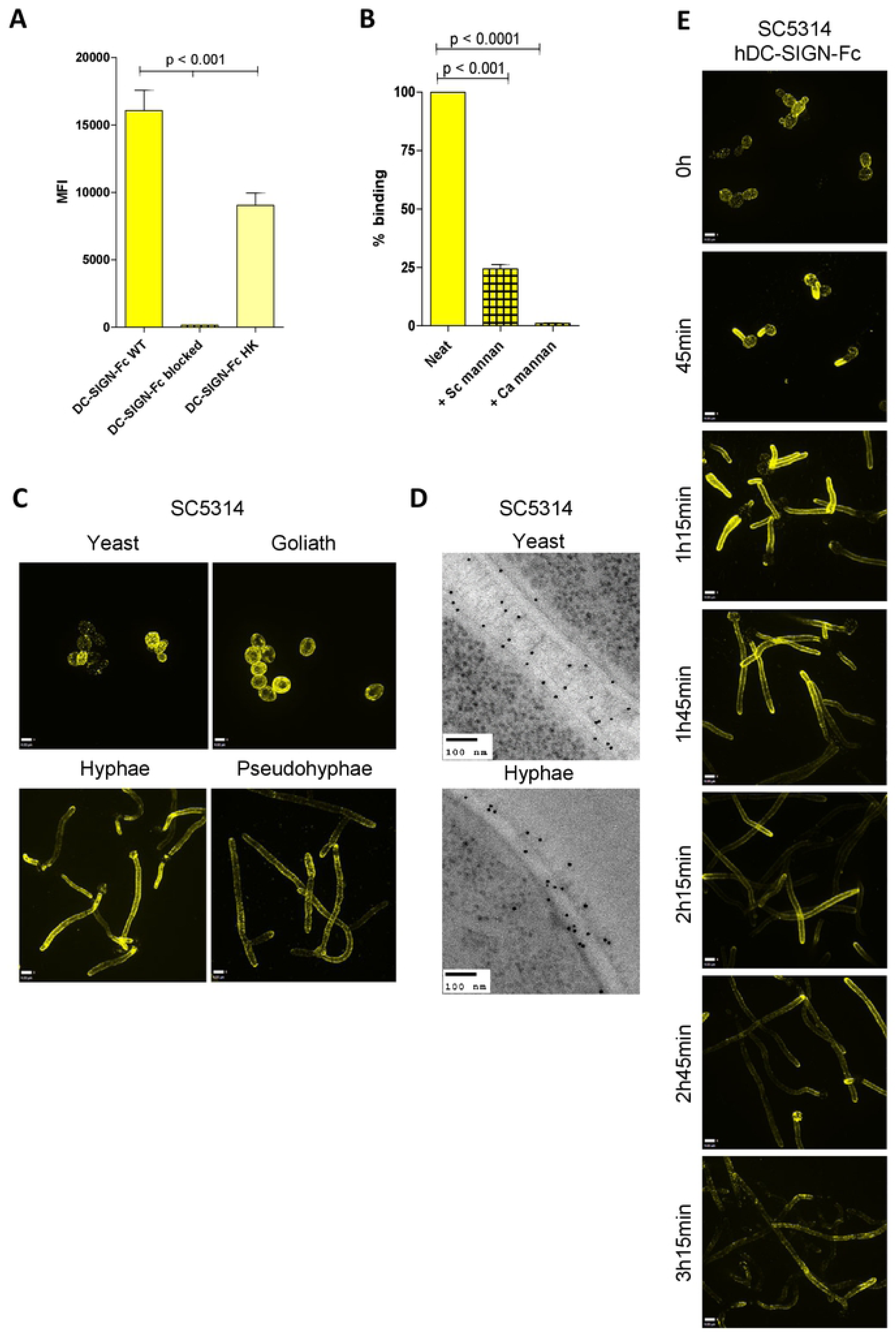
hDC-SIGN-Fc binding across the inner and outer cell wall of *C. albicans*. (A) hDC-SIGN-Fc binding to WT 4% paraformaldehyde fixed *C. albicans* (CAI4-CIp10) yeast cells compared to 65°C HK cells and hDC-SIGN-Fc blocked with *C. albicans* cell wall mannan (25 µg/ml). (B) Binding of hDC-SIGN-Fc blocked with *C. albicans* cell wall mannan (25 µg/ml) or *S. cerevisiae* cell wall mannan (25 µg/ml) to WT 4% paraformaldehyde fixed *C. albicans* yeast cells. (C) Indirect immunofluorescence staining of different *C. albicans* (SC5314, BWP17 + Clp30 for goliath cells) morphological forms. (D) Immunogold localisation of hDC-SIGN-Fc on *C. albicans* (SC5314) yeast and hyphae cells. (E) Representative images of immunofluorescence staining of hDC-SIGN-Fc binding to *C. albicans* (SC5314) hyphae over prolonged periods of growth. Data were obtained using UltraView® VoX confocal spinning disk microscope, a BD Fortessa flow cytometer and JEM-1400 Plus using an AMT UltraVUE camera for TEM images. Scale bars represent 4 µm for immunofluorescence images and 100 nm for TEMs. For immunofluorescence staining and flow cytometry experiments, 2.5 × 10^6^ cells were used per treatment. Median fluorescence values were used for quantification of hDC-SIGN-Fc binding and represented data are means ± SEM, n = 3. Statistical analyses used One Way Anova with Tukey’s post-hoc test.

### Chemical nature of C-type lectin receptor targets in fungal cell walls

To assess the nature of the targets of the CTL receptors, the binding of the Fc-lectin probes was examined using a collection of cell wall mutants including isogenic nulls with truncations in *N-* and *O-*mannans (Fig. 9). In general, mannosylation mutants exhibited marked changes in Fc-lectin recognition. The *C. albicans mnn4*Δ mutant lacks phosphomannosyl residues and consequently these mutants have uncharged cell walls [65]. The *mnn4*Δ demonstrated increased binding of dectin-2-Fc, reduced binding of CRD4-7-Fc and a similar binding profile of hDC-SIGN-Fc compared to SC5314 (Fig. 9 A). The *N-*mannan outer chain mutant *mnn2-26*Δ that lacks *N*-mannan side chains demonstrated a slight increase in dectin-2-Fc, reduced CRD4-7-Fc and a similar binding profile of hDC-SIGN-Fc compared to SC5314 (Fig. 9 A) [66]. An *N*-mannan mutant *och1*Δ, with no outer *N*-linked mannan chains and a markedly reduced phosphomannan content but increased chitin and glucan [67], resulted in much higher binding of dectin-2-Fc, CRD4-7-Fc and a dense binding pattern all over the yeast cell of hDC-SIGN-Fc compared to SC5314 (Fig. 9 A). Increased binding of some *N*-mannan Fc-lectins to mannosylation mutants can be explained by the compensatory synthesis of PRR-binding epitopes as a consequence of the mutation [67]. A *pmr1*Δ mutant with reduced phosphomannan, *O-*mannan and *N*-mannan [68] demonstrated marginally increased recognition by dectin-2-Fc and reduction of CRD4-7-Fc and similar binding of hDC-SIGN-Fc compared to wild type (Fig. 9 A). This is compatible with previous observations that dectin-2-Fc recognised inner cell wall mannans whilst CRD4-7-Fc labelled the outer mannoprotein fibrillar layer (Fig. 6 A, B). Dectin-1-Fc was used as a control and demonstrated increased binding in all backgrounds deficient in *N-* and *O-*mannan attributable to the increased exposure of inner cell wall component β-glucan (Fig. 9 A). Therefore, the cell wall glycosylation status had a major impact on the ability of CTL probes to bind *C. albicans*.

**Figure 9.**
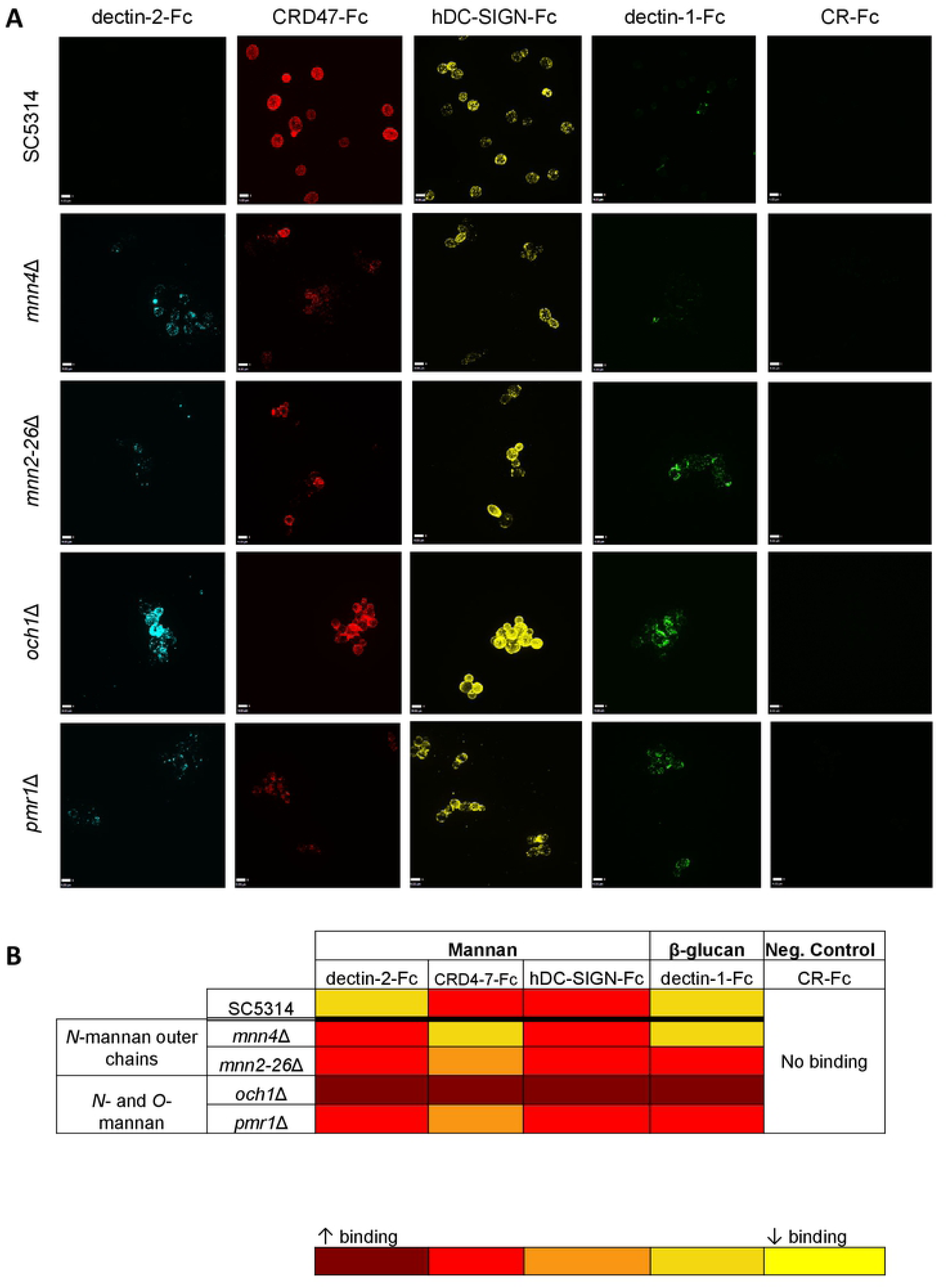
Fc-lectin binding profiles to *C. albicans* cell wall glycosylation mutants. (A) Indirect immunofluorescence images of Fc-conjugated CTL probes binding to *C. albicans O-*mannan and *N-*mannan cell wall mutants of *C. albicans* (as described in the text). (B) Heat-map of increased or reduced binding of Fc-lectins to *C. albicans* cell wall mutants relative to the SC5314 clinical isolate. Dark red (increased binding) to yellow (reduced binding). Data were obtained using UltraView® VoX confocal spinning disk microscope. Scale bars represent 4 µm.

To gain further insight into the carbohydrate recognition by the CTL receptors, we analysed these initially using a microarray comprised mostly of fungal-type saccharides and compared their binding profiles (Fig. 10, S Table 2). Both CRD4-7-Fc and hDC-SIGN-Fc showed strong binding to the *C. albicans N*-mannoprotein that is characterised by an α-1,6-mannose backbone with oligomeric α-1,2-, α-1,3-, and β-1,2-manno-oligosaccharide branches (Fig. 10 A, B, position 13, S Table 2). The two proteins bound also to other mannan-related saccharides from *S. cerevisiae* and *M. tuberculosis*, that share an α1,6-mannose backbone with α-1,2-manno-oligosaccharide branches (Fig. 10 A, B, S Table 2). In contrast, no binding was detected with dectin-2-Fc to any of Manα-1,2-Man-containing polysaccharides in the conditions of the microarray analysis, which suggests it may have less capacity to bind α-mannans of this type compared to CRD4-7-Fc and hDC-SIGN-Fc (data not shown). Dectin-1-Fc showed, as predicted, strong and highly specific binding to glucans with a β-1,3-glucosyl backbone (Fig. 10 C, S Table 2).

**Figure 10.**
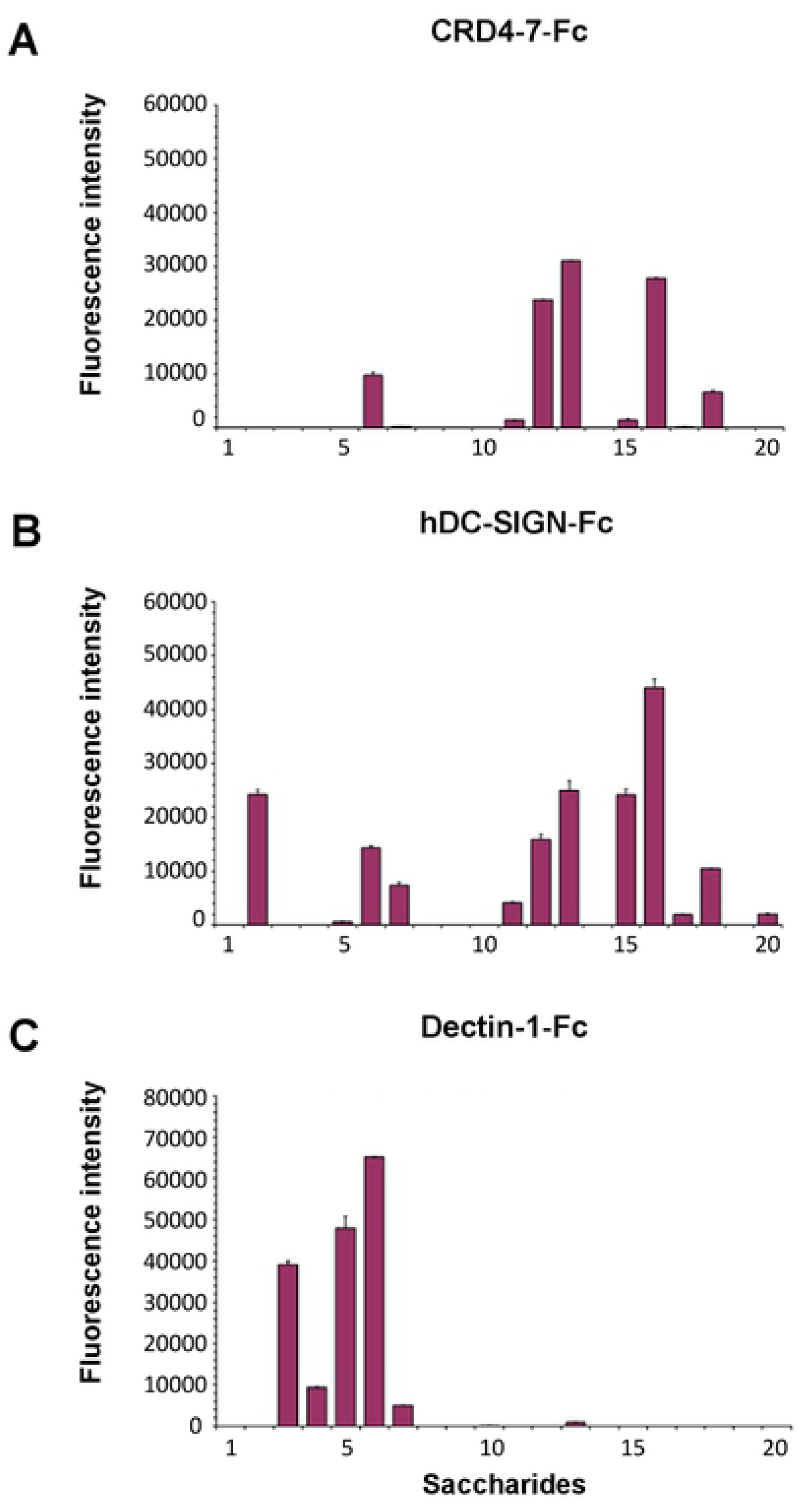
Glycan microarray analyses of Fc-lectins using the fungal and bacterial polysaccharides. Binding signals of CRD4-7-Fc (A), hDC-SIGN-Fc (B) and dectin-1-Fc (C) to a variety of fungal and bacterial polysaccharides. The saccharide positions and predominant oligosaccharide sequences are specified in the supplementary materials (S Table 2). The binding signals are means of the fluorescence intensities of duplicate spots printed at the high level of probe arrayed 0.1 ng/spot (saccharide positions 1-19) and at 5 fmol/spot (position 20). The error bars represent half the difference between the two values.

Glycan microarrays of 474 sequence-defined oligosaccharide probes (S Table 3 B) were also applied to compare the binding specificities of dectin-2-Fc, CRD4-7-Fc and hDC-SIGN-Fc. The signal intensities observed with dectin-2-Fc were the lowest overall among the three Fc probes. Dectin-2-Fc binding was detected to Man_9_GN_2_ derived probes that resemble the core *N*-mannan structures within the inner *C. albicans* cell wall (Fig. 11 A, S Table 3 A); this relatively weak and restricted binding is in agreement with previous glycan array studies [37, 69]. No binding of dectin-2-Fc was detected to oligo-mannose sequences smaller than Man_7_GN_2_. Additionally, binding was detected of dectin-2-Fc to 3’sialyl LNFPIII and a number sulphated glycans as with hDC-SIGN-Fc.

**Figure 11.**
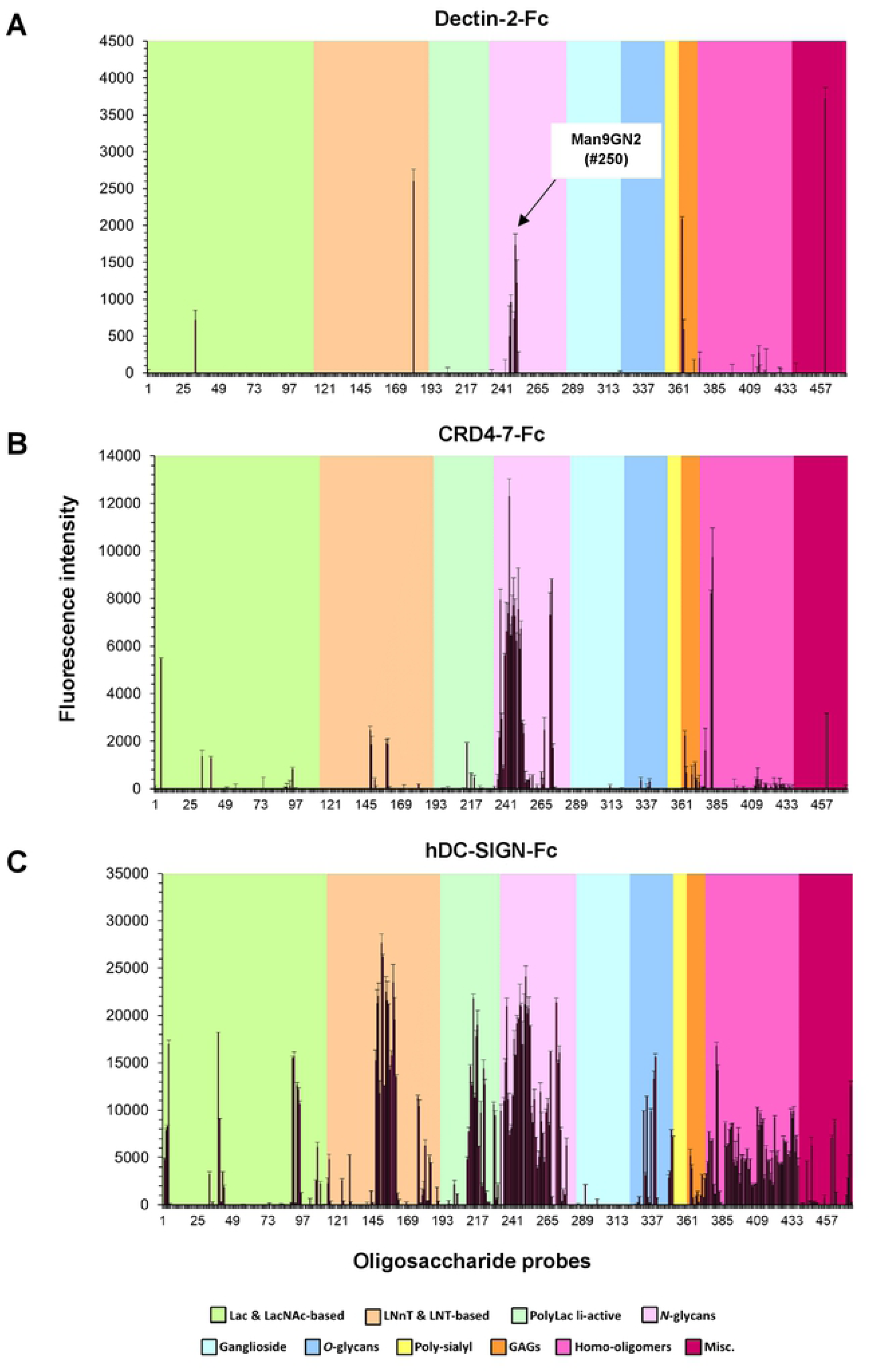
Microarray analysis of Fc-lectins using a screening array of sequence-defined glycan probes. Binding signals of dectin-2-Fc (A), CRD4-7-Fc (B) and hDC-SIGN-Fc (C) to a variety of sequence-defined polysaccharides. The glycan resembling the core *N*-mannan structure in *C. albicans* cell wall (position 250) is highlighted in dectin-2-Fc panel. The binding signals are means of the fluorescence intensities of duplicate spots printed at 5 fmol/probe. The error bars represent half of the difference between the two values. The glycan probes are grouped according to their backbone-type sequences as annotated by the colored panels: disaccharide based: lactose (Lac) and *N*-acetyllactosamine (LacNAc); tetrasaccharide based: lacto-*N*-neo-tetraose (LNnT) and lacto-*N*-tetraose (LNT); poly-*N*-acetyllactosamine (PolyLacNAc); *N*-glycans; gangliosides; *O*-glycan-related; polysialyl; glycosaminoglycans (GAGs); homo-oligomers of glucose and of other monosaccharides, and other non-classified sequences (miscellaneous, Misc). The list of glycan probes and their sequences are in supplementary materials (S Table 3 B).

CRD4-7-Fc showed mannose-related binding of high intensity to oligo/high-mannose *N*-glycan sequences, fucosylated probes including Fuc-GlcNAc and Man3FGN2, as well as β-1,4-oligomannoses (Fig. 11 B, S Table 3 A). hDC-SIGN-Fc gave strong binding signals with *N*-acetylglucosamine containing sequences including chitin-derived glycans (Fig. 11 C, S Table 3 A) and also to glucan oligosaccharide sequences with differing glucosyl linkages as was also observed in recent studies [70, 71] (Fig. 11 C, S Table 3 A)]. hDC-SIGN-Fc also gave binding to a broad range of *N*-glycans including oligo/high-mannose sequences having α-1,2-, α-1,6-and α-1,3/1,6-linked mannose, high-mannose sequences capped by Glc residues and a number of *N*-acetylglucosamine-terminating *N*-glycans; binding was also detected to β4-linked mannose oligosaccharides as with CRD4-7-Fc. Collectively, glycan array binding results are consistent with the dectin-2-Fc ligand being based on Man_9_GN_2_ found in the core *N*-mannan triantennary structure within the *C. albicans* inner cell wall, whereas CRD4-7-Fc and hDC-SIGN-Fc have a broader binding profile compared to dectin-2-Fc and can recognise oligo-mannose structures terminating in α-1,3 and α-1,6-mannose resembling to some extent those found in the outer-chain mannan structures. This is compatible with the binding patterns observed using colloidal-gold TEM.

## Discussion

CTL receptors provide a first line defence against fungal pathogens and orchestrate both innate and adaptive immunity through the recognition of fungal PAMPs. A large number of CTL receptors have been proposed to bind fungal cell wall epitopes such as mannans, β-1,3-glucan and chitin [34, 37, 47, 50, 52]. Previous studies suggested dectin-2 to recognise high mannose residues that are present on fungal cell surfaces while MR was proposed to bind branched *N*-mannan structures, with fucose, *N*-acetylglucosamine sugar residues and mannose-capped lipoarabinomannan (ManLAM) on the mycobacterial cell wall [34, 37, 72]. hDC-SIGN has been demonstrated to recognise galactomannan, mannose- and fucose-containing glycoconjugates and *N*-linked mannose rich components in *C. albicans* cell wall [38, 73]. There is however limited knowledge of the structural arrangement, spatial distribution and variation in mannoside architecture of the cell wall of fungal pathogens, which this study addresses. We deployed recombinant CTL-Fc proteins as probes to explore the distribution, regulation and chemical structure of fungal ligands for these receptors within the cell wall, in particular for dectin-2, MR and hDC-SIGN. We demonstrated that there is considerable intra- and inter-species variability in the expression of key mannan epitopes. We mapped the patterns of binding to these ligands during growth and cellular morphogenesis and observed marked spatial segregation and an unexpected clustering of certain mannan epitopes.

The analysis of *C. albicans* cell wall mutants corroborated the occurrence of inner cell wall epitopes for dectin-2-Fc and superficial mannan ligands for MR and hDC-SIGN-Fc. An *och1*Δ mutant, which has a defect in its ability to synthesise outer chain *N*-glycans, had been shown previously to have exposed inner cell wall components [67]. This mutant also exhibited increased binding by dectin-2-Fc and has been shown to have an elaborated α-1,2- and α-1,3-mannan side chains to the Man_8_GlcNAc_2_ core triantennary complex which is a ligand that promotes CRD4-7-Fc and hDC-SIGN-Fc binding. Other mutants with decreased *N*-mannan outer chains, including *mnn2-26*Δ, *pmr1*Δ [66, 68] bound less CRD4-7-Fc and hDC-SIGN-Fc and more dectin-2-Fc commensurate with an increased exposure of the inner wall layers. Moreover, the *mnn4*Δ mutant, which lacks cell wall phosphomannan that confers a negative charge on the wall [65] had reduced binding by CRD4-7-Fc and hDC-SIGN-Fc but increased dectin-2-Fc affinity. These observations complement previous studies suggesting that cell surface glycosylation profoundly influences the efficiency of pathogen recognition by immune receptors [9, 11, 13, 23, 74]. Glycan microarray data with the Fc-lectins and the fungal-type saccharides are consistent with CRD4-7-Fc and hDC-SIGN-Fc but not dectin-2-Fc recognizing α-1,6-Man backbone with oligomeric α-1,2-, α-1,3-, and β-1,2-Man branches which comprise *C. albicans* outer wall *N*-mannan branches. This contrasts with the weak binding detected of dectin-2-Fc to the high-mannose Man_9_GN_2_ probe with terminal Manα-1,2-Man sequences, which resembles *C. albicans* core *N*-mannan within the inner cell wall. This is in agreement with published data that showed the Manα-1,2-Man sequence on the Man_9_GN_2_ to be a primary target for dectin-2 binding [37, 69], and that this receptor could adopt a geometry of the binding site that accommodates the internal residues of the Manα-1,2-Manα-1,2-Man-(D1 branch) and Manα-1,2-Manα-1,3-Man-(D2 branch) trisaccharide sequences [37, 69]. This prompts a hypothesis that dectin-2 binds to internal sequences in fungal mannan polysaccharides, expressing higher density of the ligands and achieving multivalent binding.

We examined the immunological signature of clinically-relevant *Candida* species and *S. cerevisiae* and demonstrated that there was no simple correlation between phylogenetic relationships between fungal species and CTL epitope distribution. Despite *C. albicans* and *C. dubliniensis* sharing approximately 95% genome identity [75] the patterns of CTL probe binding contrasted markedly. The immunologically relevant mannan structures are produced by activities of multiple families of mannosyltransferases, of which genetic variation and regulation may not map simply to phylogenetic relationships. We focused mainly on *C. albicans* as the model organism for further mapping of cell wall dynamics. Previous studies demonstrated that dectin-1 recognition of *C. albicans* was influenced by fungal strain type because chitin levels modulated the accessibility of dectin-1 ligands [76]. Analogously, binding of mannan-recognising CTL probes varied in different *C. albicans* strains and species. Previous studies have identified virulent and attenuated isolates of this organism [58, 59], however, Fc-lectin probes recognised both virulent and avirulent strains with similar binding patterns and there was no simple correlation between virulence and CTL probe binding (Fig. 2 A). Previous studies have also demonstrated a strong correlation between the capacity to form hyphae and virulence [77–81]. The intensity and distribution of mannan-specific CTL binding to hyphae was found to vary during hypha elongation. It is possible that the efficiency by which invading hyphae induce or escape immune surveillance ultimately influences virulence.

We also demonstrated that *C. albicans* growth phase influenced expression and exposure of cell wall components. Batch growth of *C. albicans* yeast culture revealed increased dectin-1-Fc binding during the exponential growth phase. This is likely to be due to increased β-glucan exposure and number of bud scars on actively dividing cells. As batch culture of cells transitioned into the stationary phase, a reduction in dectin-1-Fc and increased dectin-2-Fc binding was observed that was likely to be related both to the degree of mannan shielding of β-glucan as the cell wall was reorganised. CRD4-7-Fc demonstrated similar binding patterns during all growth phases of the yeast batch culture, highlighting the diversity in availability of the cell wall manno-oligosaccharide sequences and suggesting that the CRD4-7-Fc (mannose receptor) ligand is superficial in the cell wall. This suggestion was supported by colloidal gold-TEM images showing CTL binding patterns at the ultrastructural level, and is in accord with a previous study which demonstrated that MR was not required for host defence in a systemic candidiasis mouse model [82].

The hyphal cell wall is also modified during the process of hyphal extension [17, 61]. In this study we mapped the distribution of CTL epitopes on hyphal cell walls over a period of prolonged hyphal growth and demonstrated that dectin-2-Fc and CRD4-7-Fc ligands were densely concentrated at the emerging germ tube apex. Previous studies on *C. albicans* suggested that mannan-recognising dectin-2 is a hypha-specific CTL [49], however, we demonstrate here that the distribution and expression of dectin-2 epitopes on yeast cells varies between different *Candida* species. There was little availability of dectin-1-Fc ligands and by inference β-1-3 glucan on hyphae, possibly because of the lack of bud scars on hyphae as discussed below. This supports previous studies describing attenuated dectin-1 activation by hyphal cells [63]. As hyphae elongated, binding of mannan-recognising probes dectin-2-Fc and CRD4-7-Fc was reduced and became more punctate while binding of dectin-1-Fc to lateral cell walls increased. This finding supports observations that have demonstrated that β-glucan becomes exposed late during disseminated *C. albicans* infection [61]. Interestingly, binding of CRD4-7-Fc to *C. albicans* germinating cells demonstrated a reduction in mother yeast cell recognition as the germ tube elongated. It is possible that the mannosylated structures on yeast cell mannoproteins are chemically modified during hyphal elongation so they are less able to be recognised by the MR. However, multiple mannan-detecting PRRs were able to bind each of the various cell types of *C. albicans* that were studied. This redundancy in CTL engagement presumably confers advantage in recognising fungal pathogens whose surfaces and shapes are remodelled in response to host invasion and colonisation of different microenvironments.

Morphological alterations of *C. albicans*, and other fungal pathogens, is often correlated with their ability to thrive within the human host [83]. These cellular transitions are coupled to changes in their cell wall composition which makes them a moving target for immune detection [25, 84, 85]. *C. albicans* forms cells that are more or less elongated [83]. Synchronously dividing elongated yeast cells (pseudohyphae) are considered by many to be a distinct cell form, and these cell types are common to all *Candida* species. The tested Fc-lectin probes also demonstrated intermediate binding affinities to pseudohyphae of *C. albicans* (Fig. 4), complementing a recent study describing intermediate cytokine profiles to *C. albicans* filamentous forms [86]. Goliath cells are recently recognised as an enlarged cell type of *C. albicans* that may play an important role in commensalism and persistence in the gut [62]. We observe that dectin-2-Fc weakly recognises *C. albicans* yeast cells but has increased binding on goliath cells (Fig. 4 A), potentially due to cell expansion and concomitant exposure of inner cell wall epitopes or a novel arrangement of mannoproteins on the surfaces of goliath cells that may expose more dectin-2-recognising mannan epitopes. Dectin-2-Fc also bound well to hyphae of *C. albicans*, in particular at the hyphal tip (Fig. 4 A). The cell wall of the hyphal apex is thinner and the polysaccharides are less crystalline and less cross-linked. This may facilitate access of high molecular weight PRRs to inner cell wall ligands. Both mannose receptor (CRD4-7-Fc) and hDC-SIGN-Fc probes demonstrated similar binding to different morphologies and their binding was not greatly affected by mild heat treatments suggesting their epitopes are superficial. As predicted, dectin-1-Fc binding was concentrated at the bud scars of yeast and goliath cells, but binding was punctate on hyphal cells that lack bud scars. Punctate binding patterns have been reported elsewhere, in particular for dectin-1 binding revealed by super-resolution microscopy in which binding became increasingly associated with highly granular multi-glucan surface exposures in response to caspofungin treatments [87].

Using multiple approaches, we provide evidence that individual mannan-recognising CTLs recognise different mannan structures that are displayed in distinct binding patterns. CRD4-7-Fc (MR) and hDC-SIGN-Fc recognised α-1,6-mannose backbone with oligomeric α-1,2, α-1,3, and β-1,2-mannan branches while dectin-2-Fc bound Man_9_GN_2_, which has close similarities to the core *N*-mannan structure in *C. albicans* inner cell wall. Heat-killing and immunofluorescent staining of *C. albicans* cell wall mutants as well as immunogold-labelling and TEM microscopy supported the conclusion that MR and DC-SIGN recognise outer chain *N*-mannan whilst dectin-2 recognises core mannans that are closer to the amide linkage polypeptide of the mannoprotein at the base of the fibrillar layer of the outer cell wall. We observed that CRD4-7-Fc and hDC-SIGN-Fc epitopes were diffusely organised across the plasma membrane plane and within the outer wall mannan fibrils while dectin-2-Fc epitopes were clustered within the inner cell wall in both yeast and hyphal *C. albicans* cells. Clustering of the dectin-2-Fc epitope requires further investigation but may suggest cooperative binding whereby binding of a single dectin-2-Fc could stabilize the target ligand in a way that facilitates bindings of additional PRR molecules. Alternatively certain cell wall proteins that display dectin-2 binding ligands might accumulate in microdomains within the inner cell wall. Membranes of eukaryotic cells are known to organise some proteins into specialised microdomains which compartmentalise cellular processes and serve as organising centres which assemble signalling molecules, facilitate protein and receptor trafficking, vesicular transport and signalling events [88, 89]. Such microdomains have also been described in bacterial membranes and a recently published study suggested that similar microdomains might also exist in fungal cell walls [90]. Therefore, clustered dectin-2-Fc binding could be related to the accumulation of specialised glycosylated proteins at certain parts of the inner cell wall. Nevertheless, these attractive hypothesis should be addressed by future studies.

In conclusion, we have demonstrated that mannan epitopes are differentially distributed in the inner and outer layers of fungal cell wall in a clustered or diffuse manner. Immune reactivity of fungal cell surfaces was not correlated with relatedness of the fungal species, and mannan-detecting receptor-probes discriminated between cell surface components generated by the same fungus growing under different conditions. Moreover, individual mannan-recognising CTL probes conferred specificity for distinct mannan epitopes. These findings indicate that the fungal cell wall structures are highly structured but dynamic, and that immune recognition is likely to involve PRRs acting alone and in concert to mount effective immune responses. This type of CTL ligand variation carries significant impact on the design of future fungal diagnostics, vaccine and therapeutics.

## Materials and Methods

### Expression and purification of Fc conjugated C-type Lectin Receptor proteins

HEK293T cells stably expressing murine dectin-1-Fc and dectin-2-Fc fusion proteins were cultured as described previously [36, 37]. Briefly, cells were cultured in T175 flasks and supernatants were collected. Zeocin (0.4 mg/ml) (Thermo Scientific) was used for selection of cells expressing Fc conjugated proteins. Large scale transient transfections for murine CRD4-7-Fc and CR-Fc were carried out using 100 µg of total plasmid DNA and 100 ml of suspension cultured Expi293F cells (Life Technologies). Supernatants were harvested at day 6. Fc conjugated protein concentrations in supernatants were quantified by ELISA. ELISA plates (Thermofisher) were coated with 100 µl of supernatants and incubated overnight at 4°C. Plates were washed with PBS + Tween 20 (0.05 %) and blocked with 200 µl of 10 % FCS in 1 X PBS for 2 h. Plates were washed with 1 X PBS + Tween 20 (0.05 %) and 100 µl of secondary anti-human antibody conjugated to HRP (Jackson Immunoresearch) diluted 1/10000 in PBS was added and incubated at room temperature (RT) for 1 h. Wells were washed with 1 X PBS + Tween 20 (0.05 %) and 100 µ of TMB (Thermofisher) were added to develop. Reaction was stopped with 50 µl 2N H_2_SO_4_ and plates were assayed on a spectrophotometer at 450_nm_ with the necessary λ correction at 570_nm_. Fc chimeric proteins were purified via affinity based Fast Protein Liquid chromatography using Prosep® Ultra resin (Millipore). Fc conjugated proteins were eluted with 0.1 M glycine (pH 2.5) before neutralisation with 1 M Tris buffer (pH 8) and then dialysed in 1 X PBS overnight. Fc conjugated protein concentration was quantified using NanoVue Spectrophotometer (GE Healthcare).

### QC of purified Fc-lectins

Purified proteins were checked via SDS-PAGE gel analysis using 4-12 % Bis-Tris SDS-PAGE gels under reducing conditions (S Fig. 1). ELISA was carried out for confirmation of binding to original target using ELISA protocol described above (S Fig. 1). ELISA plates were coated with live *C. albicans* yeast cells, 25 µg/ml *S. cerevisiae* mannan (SIGMA), 100 µg/ml *C. albicans* yeast β-glucan or PBS overnight (S Fig. 1). Fc chimeric proteins were added at 5 µg/ml and serial doubling dilutions were performed to confirm concentration-dependent binding.

### Comparison of fungal strains under different parameters

For comparison of fixed and heat-killed *C. albicans* (CAI4-CIp10) cells, a single colony was inoculated into 10 ml YPD (1% yeast extract, 2% glucose, 2% peptone) and incubated overnight at 30°C, 200 rpm. Overnight culture was washed in 1 X PBS and 2.5 × 10^6^ cells were either fixed with 4 % paraformaldehyde or kept at 65°C in a heat block for 2 h prior to staining. For comparison of *C. albicans* (CAI4-CIp10) culture overtime, OD_600_ of overnight culture was measured and culture was diluted to OD_600_ of 0.1 in 50 ml YPD in 250 ml flasks. Cells were collected at OD_600_ 0.2, 0.4, 0.6, 1 and 18, washed in 1 X PBS, fixed with 4 % paraformaldehyde at RT for 45 min, washed and then stained. For comparison of different *C. albicans* isolates and cell wall mutants, cells were fixed at OD_600_ ~ 0.5 prior to staining (S Table 1). For comparison of different *Candida* species and *S. cerevisiae*, cells were fixed in stationary phase, OD_600_ ~ 18 (S Table 1). Samples were stained as described below and analysed on BD Fortessa flow cytometer or in 3D on an UltraVIEW® VoX spinning disk confocal microscope. Three independent biological replicates were performed per sample.

### Conditions for generating different morphologies of *C. albicans*

Single colonies of *C. albicans* were inoculated into 10 ml YPD and incubated overnight at 30°C, 200 rpm. To induce hypha formation, cultures were diluted 1:1333 in milliQ water and then adhered on a poly-L-lysine coated glass slide (Thermo Scientific, Menzel-Gläser) for 30 min prior to incubation in pre-warmed RPMI + 10 % FCS at 37°C for 45 min- 3 h 15 min depending on the tested parameter. Slides were than washed in DPBS and fixed with 4 % paraformaldehyde. *C. albicans* pseudohyphae were produced using published conditions with modifications [91]. Overnight culture was collected by centrifugation, washed twice with 0.15 M NaCl, resuspended in 0.15 M NaCl and incubated at room temperature for 24 h to induce starvation. After 24 h, cells were transferred into RPMI 1640 at a concentration of 1×10^6^ and incubated at 30°C 200rpm for 6 h prior to fixation with 4 % paraformaldehyde. Fixed cells were stained and imaged as described below. To induce goliath cell formation, a single colony of *C. albicans* was inoculated in 4 ml SD media (2% glucose, 6.7g/L yeast nitrogen base without amino acids) and incubated for 24 h at 30°C 200 rpm [62]. Following incubation, 600 µl of culture were washed in three times milliQ water and resuspended in 600 µl milliQ water prior to OD_600_ measurement. To elicit zinc starvation, and hence goliath cell formation, washed cells were inoculated into 4 ml of Limited Zinc Medium (LZM) at OD_600_ 0.2. LZM culture was incubated for 3 days prior to fixation with 4% paraformaldehyde and staining [62].

### Immunofluorescent staining of Fc-lectins binding to fungal cells

Yeast cells were counted using an Improved Neubauer haemocytometer and 2.5 × 10^6^ cells were transferred into V-bottomed 96-well tissue culture plates. Plates were centrifuged at 4000 rpm 5 min and supernatants were removed. Samples of 1 µg/ml dectin-1-Fc in PBS, 1 % (v/v) FCS or 2 µg/ml of dectin-2-Fc, CRD4-7-Fc, CR-Fc or DC-SIGN-Fc (Thermofisher) in binding buffer (BB) (150 mM NaCl, 10 mM Tris pH 7.4, 10 mM CaCl_2_ in sterile water + 1% FCS) were then added to appropriate wells and incubated for 45 min on ice. Cells were washed once in PBS + 1% FCS for dectin-1-Fc or BB buffer for dectin-2-Fc, CRD4-7-Fc, CR-Fc and DC-SIGN-Fc and then stained with Alexa Fluor® 488 goat anti-human IgG antibody (Life Technologies) diluted 1/200 in PBS + 1% FCS or BB buffer and incubated 30 min on ice. Stained cells were washed twice before final resuspension in PBS + 1% FCS or BB buffer. For staining filamentous cells, Fc protein in PBS + 1% FCS or BB buffer at the same concentration as above was added on top of poly-L-lysine slides. Samples were analysed using a BD Fortessa flow cytometer where 10000 events were recorded for each sample from three independent experiments. Median fluorescence intensity for asymmetric peaks and mean fluorescence intensity for symmetric peaks was calculated for each sample using FlowJo v.10 software. Alternatively, 5 µl of yeast cells were added on a poly-L-lysine coated glass slides (Thermo Scientific, Menzel-Gläser) prior to imaging in 3D on an UltraVIEW® VoX spinning disk confocal microscope (Nikon, Surrey, UK).

### High Pressure Freezing (HPF) of samples for immunogold labelling of *C. albicans* with Fc-lectins for Transmission Electron Microscopy (TEM)

Yeast and hyphal *C. albicans* samples were prepared using high-pressure freezing by EMPACT2 high-pressure freezer and rapid transport system (Leica Microsystems Ltd., Milton Keynes, United Kingdom). Cells were freeze-substituted in 1% acetone (w/v) OsO_4_ by using a Leica EMAFS2 prior to embedding in Spurr’s resin and polymerizing at 60°C for 48 h. Ultrathin sections were cut using a Diatome diamond knife on a Leica UC6 ultramicrotome and sections were mounted onto formvar coated copper grids. Subsequently, sections on formvar coated copper grids were blocked in blocking buffer (PBS + 1% (w/v) BSA and 0.5% (v/v) Tween20) for 20 min prior to incubation in three washes in binding buffer (150 mM NaCl, 10 mM Tris pH 7.4, 10 mM CaCl_2_ in sterile water, 1% FCS) for 5 min. Sections were then incubated with Fc chimeric proteins (5 µg/ml for yeast and 10 µg/ml for hyphae) for 90 min before six washes in binding buffer for 5 min. Fc protein binding was detected by incubation with Protein A conjugated to 10 nm gold (Aurion) (diluted 1:40 in PBS + 0.1% (w/v) BSA) for 60 min prior to six 5 min washes in PBS + 0.1% (w/v) BSA followed by three, 5 min washes in PBS, and three, 5 min washes in water. Sections were stained with uranyl acetate for 1 min prior to three 2 min washes in water and were left to dry. TEM images were taken using a JEM-1400 Plus using an AMT UltraVUE camera.

### Glycan microarray analyses of Fc-lectins

The binding specificities of the Fc-lectin receptors were analysed using two types of carbohydrate microarrays: (1) a microarray designated ‘Fungal and Bacterial Polysaccharide Array’ featuring 19 saccharides (polysaccharides or glycoproteins) and one lipid-linked neoglycolipid (NGL) derived from the chitin hexasaccharide (S Table 2); and (2) a screening microarray of 474 sequence-defined lipid-linked glycan probes, of mammalian and non-mammalian type (S Table 3 B) essentially as previously described [92]; these probes are a subset of a recently generated large screening microarray containing around 900 glycan probes (in-house designation “Array Sets 42-56”, which will be published in detail elsewhere).

The Fc-lectin binding was performed in both types of arrays essentially as described [28]. In brief, after blocking the slides with 0.02% v/v Casein (Pierce) and 1% BSA (Sigma) diluted in HBS (10 mM HEPES-buffered saline, pH 7.4, 150 mM NaCl) with 10 mM CaCl_2_, the microarrays were overlaid with the Fc-lectins precomplexed with the biotinylated goat anti-human IgG (Vector) for 2 hours. The Fc-lectin-antibody complexes were prepared by preincubating the –Fc-lectin with the antibody at equimolar ratios for 1 hour, followed by dilution in the blocking solution to give the final Fc-lectin concentration: dectin-2-Fc 10 µg/ml, CRD4-7-Fc 20 µg/ml and hDC-SIGN-Fc 2 µg/ml. Dectin-1-Fc used as control for the Fungal and Bacterial Array was analysed non-precomplexed at 20 µg/ml in the blocking solution 0.5% v/v casein (Pierce) in HBS. The binding was detected with Alexa Fluor-647-labeled streptavidin (Molecular Probes, 1 μg/ml). All steps were carried out at ambient temperature. Details of the glycan library including the sources of saccharides, the generation of the microarrays, imaging, and data analysis are in the Supplementary glycan microarray document (S Table 4) in accordance with the Minimum Information Required for A Glycomics Experiment (MIRAGE) guidelines for reporting glycan microarray-based data [93].

## Acknowledgements

We thank Luisa Martinez-Pomares and Darryl Jackson from Nottingham University for providing CRD4-7-Fc and CR-Fc plasmids; Fiona M. Rudkin for advice on protein expression and purification; Louise Walker for high pressure freezing of samples for TEM analysis; David Williams for *C. albicans* purified mannan and β-glucan preparations; Dhara Malavia for providing *C. albicans* goliath cells, the University of Aberdeen Core Microscopy & Histology Facility (Kevin MacKenzie, Lucinda Wight, Debbie Wilkinson), Iain Fraser Cytometry Centre (Raif Yuecel). The neoglycolipid-based glycan microarrays contain several saccharides provided by collaborators whom we thank as well as members of the Glycosciences Laboratory for their collaboration in the establishment of the microarray system.

## Author Contributions

IV, JAW, GDB & NARG conceived and designed the experiments; IV, LMS, MHTS, ASP performed experiments; IV, JAW, LMS, ASP, YL & NARG analysed data and NARG, GDB, WC & TF contributed reagents / materials/ analysis tools: IV, LMS, ASP & NARG wrote the paper and all authors revised and commented on the manuscript.

---

**S Figure 1.**
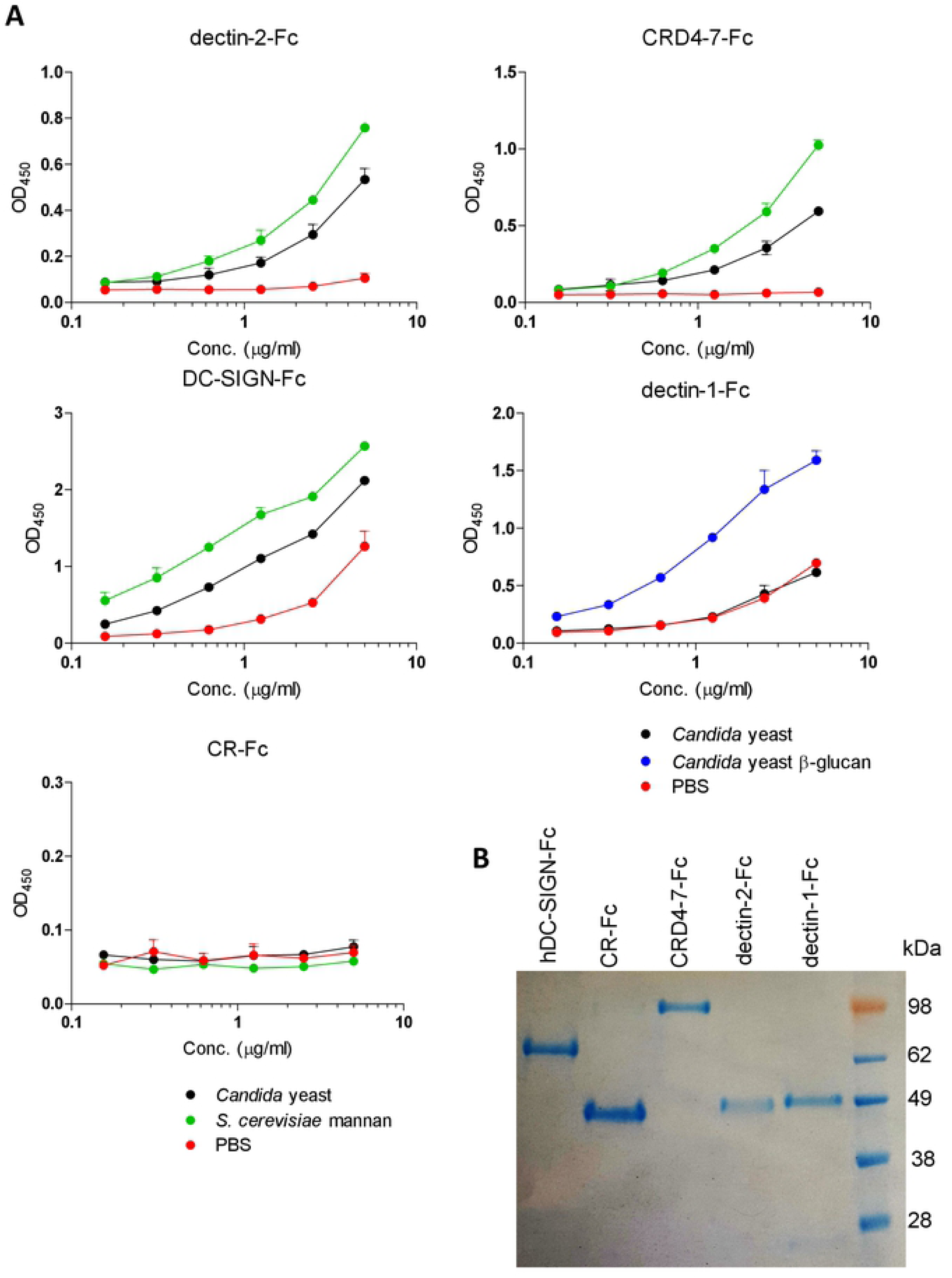
Concentration response curves of Fc-lectin binding to target antigens via ELISA. Purified Fc-lectin probes were screened against whole *C. albicans* (SC5314) yeast cells (black), purified cell wall mannan (green) or purified yeast β-glucan (blue) (A). Fc-lectin integrity was checked via reducing SDS-Page, expected band sizes were dectin-1-Fc 55 kDa, dectin-2-Fc 55 kDa, CRD4-7-Fc 110 kDa, CR-Fc 50 kDa, hDC-SIGN-Fc 69kDa (B).

